# Versatile Cell Penetrating Peptide for Multimodal CRISPR Gene Editing in Primary Stem Cells

**DOI:** 10.1101/2024.09.23.614499

**Authors:** Josh P. Graham, Jose Gabriel Castro, Lisette C. Werba, Luke C. Fardone, Kevin P. Francis, Anand Ramamurthi, Michael Layden, Helen O. McCarthy, Tomas Gonzalez-Fernandez

## Abstract

CRISPR gene editing offers unprecedented genomic and transcriptomic control for precise regulation of cell function and phenotype. However, delivering the necessary CRISPR components to therapeutically relevant cell types without cytotoxicity or unexpected side effects remains challenging. Viral vectors risk genomic integration and immunogenicity while non-viral delivery systems are challenging to adapt to different CRISPR cargos, and many are highly cytotoxic. The arginine-alanine-leucine-alanine (RALA) cell penetrating peptide is an amphiphilic peptide that self-assembles into nanoparticles through electrostatic interactions with negatively charged molecules before delivering them across the cell membrane. This system has been used to deliver DNAs, RNAs, and small anionic molecules to primary cells with lower cytotoxicity compared to alternative non-viral approaches. Given the low cytotoxicity, versatility, and competitive transfection rates of RALA, we aimed to establish this peptide as a new CRISPR delivery system in a wide range of molecular formats across different editing modalities. We report that RALA was able to effectively encapsulate and deliver CRISPR in DNA, RNA, and ribonucleic protein (RNP) formats to primary mesenchymal stem cells (MSCs). Comparisons between RALA and commercially available reagents revealed superior cell viability leading to higher numbers of transfected cells and the maintenance of cell proliferative capacity. We then used the RALA peptide for the knock-in and knock-out of reporter genes into the MSC genome as well as for the transcriptional activation of therapeutically relevant genes. In summary, we establish RALA as a powerful tool for safer and effective delivery of CRISPR machinery in multiple cargo formats for a wide range of gene editing strategies.

## 1. Introduction

Clustered regularly interspaced short palindromic repeats (CRISPR) is a precise and tunable tool for reliable genomic and transcriptomic regulation [1]. This technology is based on the use of CRISPR associated (Cas) endonucleases which are directed by guide RNAs (gRNAs) to specific genomic loci to facilitate the knock-out, knock-in, activation, or inhibition, of specific genes [2, 3, 4]. Compared to previous gene editing systems such as zinc finger nucleases (ZFNs) and transcription activator-like effector nucleases (TALENS), CRISPR adds ease of design, increased specificity and efficiency, and multiplexed targeting [5]. Despite these advantages, the use of CRISPR for the editing of primary cells is limited by the inability to deliver the necessary machinery in an efficient manner while preserving cell viability and function [6]. Specifically, this is key for applications involving the editing of clinically relevant stem and progenitor cells such as mesenchymal stromal cells (MSCs). These cells must maintain multipotency, proliferation, and immunomodulation post-editing to ensure the production of a large number of healthy cells with the desired therapeutic properties [7].

Among CRISPR delivery strategies to MSCs, viral vectors are often preferred due to strong, long-lasting expression. However, viral vectors are inherently immunogenic, activating antigen presenting cells and innate immune cells causing prolonged inflammation [8, 9, 10, 11]. They are also susceptible to random genomic integration, risking constitutive Cas9 expression which induces significant toxicity in primary stem cells and increases off-target genomic mutations [12, 13]. Viral vectors can also dysregulate natural stem cell pathways [14, 15]. Adenoviral and lentiviral transduction of MSCs was reported to reduce proliferation, induce senescence, and impair differentiation capacity, limiting overall therapeutic potential [16, 17]. In contrast, non-viral vectors reduce the risk of random genomic insertions and can deliver larger genetic payloads with a simpler production process [18]. However, these are not exempt of negative side effects. Physical methods such as electroporation or microinjection enable high editing efficiencies but lead to high cell death [19]. Moreover, physical methods require costly laboratory equipment and are most effective for *ex vivo* cell modification [20]. In contrast, non-viral nanoparticles composed of polymers, lipids, or inorganic reagents have less drastic effects on cell viability and can be delivered *in vivo* through systemic or local injection, or via incorporation into 3D biomaterials [21, 22]. Lipid-based strategies have attracted much attention due to their use in mRNA vaccines and have been employed across multiple CRISPR applications [23, 24, 25, 26]. These strategies can achieve high transfection rates allowing for transient gene expression. Additionally, lipids are biodegradable, which prevents severe cytotoxicity and inflammation. However, many commercial lipid vectors still have cytotoxic effects on transfected cells, leading to transcriptomic dysregulation that results in cell death and the disruption of stem cell’s differentiation and proliferation capacity [27, 28, 29]. Jacobsen *et al.* demonstrated that Lipofectamine transfection alone resulted in transcriptomic changes affecting over 1,000 genes [30]. These unexpected transcriptomic effects undermine the precision of CRISPR gene editing and reduce the number of healthy cells with the desired therapeutic properties.

An additional limitation of current non-viral strategies is the lack of versatility to deliver CRISPR machinery in different molecular formats. Different CRISPR applications require CRISPR components to be delivered in specific cargo types, such as plasmid DNA (pDNA), messenger RNA (mRNA), guide RNA (gRNA), or ribonucleoprotein complexes (RNPs). The choice of CRISPR format depends on factors like the target gene, cell type, and editing goals. However, existing systems lack the capability to efficiently deliver CRISPR machinery in multiple cargo formats, hindering the scalability, cost-efficiency, and standardization of biomanufacturing processes. Each CRISPR component has distinct physiochemical properties that limit their delivery through a broad range of non-viral reagents [31]. For example, Walther *et al.* attempted to deliver CRISPR components in both mRNA and RNP formats using an ionizable lipid nanoparticle [32]. While this method effectively delivered mRNA, RNP delivery resulted in inefficient nuclear localization and low editing rates (<10%) [32]. Similarly, while Lipofectamine 3000 efficiently delivers CRISPR in pDNA format, it falls short for mRNA and RNP delivery, requiring the Lipofectamine MessengerMAX and CRISPRMAX reagents respectively. This reliance on multiple reagents burdens cell biomanufacturing processes; therefore, new systems are needed to safely deliver CRISPR machinery in diverse molecular formats.

Cell penetrating peptides (CPPs) are a tunable alternative for CRISPR delivery. They can be easily designed and produced and enable competitive transfection rates with reduced cytotoxicity. CPPs have been used to deliver CRISPR components through incorporation into composite lipid or polymeric nanoparticles, direct fusion to Cas proteins, or as self-assembling nanoparticles with CRISPR cargos. Incorporation of CPPs into lipid or cationic polymer nanoparticles can increase transfection rates and add cell-specificity by targeting specific cell surface receptors [33, 34, 35, 36]. However, this method does not address lipid-associated cytotoxicity and transcriptomic dysregulation [34, 35, 36]. Alternatively, CPPs can be fused to Cas endonucleases to allow cellular uptake with minimal effects on cell viability [36]. This method requires careful design since initial attempts achieved low editing efficiencies due to interference between the covalently bound CPP and the endonuclease active site or gRNA binding domains [36]. More recently, Zhang *et al.* (2024) fused Cas9 or Cas12a to the transactivator of transcription (TAT) peptide sequence [37]. This allowed for effective CRISPR RNP delivery, resulting in ∼70% editing efficiency in human CAR-T cells [37]. However, the need of a secondary peptide for endosomal escape limits the applicability of this approach *in vivo* due to challenges with ensuring that both components reach all target cells [37]. Self-assembling CPP nanoparticles can facilitate the cytoplasmic delivery of CRISPR components without these limitations. Recently, Foss *et al.* (2023) identified the A5K peptide for delivering Cas9 or Cas12a RNPs to primary human T and B cells [33]. This approach achieved 67-68% knock-out efficiency with no cytotoxicity and minimal transcriptomic perturbation [33]. However, A5K could not deliver the DNA donor sequences required for homology directed repair (HDR) and gene knock-in [33]. Adeno-associated virus (AAV) vectors were required to supplement peptide delivery, which introduces the risk of unexpected side effects such as immunogenicity and complicates manufacturing processes [33]. While existing CPP-CRISPR strategies demonstrate high knock-out efficiencies through RNP delivery, they are unable to deliver diverse CRISPR cargos. This highlights the need of new CPPs capable to effectively facilitate all CRISPR strategies without the need of additional peptides or supporting viral vectors.

The arginine-alanine-leucine-alanine (RALA) peptide is a helical, amphiphilic CPP that self-assembles into nanoparticles in the presence of negatively charged cargos due to electrostatic interactions. This simple assembly mechanism allows for complexing with DNAs, RNAs, and small anionic molecules such as tricalcium phosphate and the chemotherapy drug 5-fluorouracil [38, 39, 40, 41]. These nanoparticles primarily cross the cell membrane through clathrin-mediated endocytosis, but can also use other endocytosis pathways or directly traverse the membrane without forming an endosome [39]. This capacity for multiple transfection pathways expands RALAs applicability to hard-to-transfect cell types including primary MSCs, dendritic cells, dermal fibroblasts, and nucleus pulposus cells [7, 41, 42, 43, 44, 45, 46, 47]. Furthermore, RALA achieves this with minimal cytotoxicity, consistently preserving cell viability compared to lipid-based methods for DNA and RNA delivery [39, 48]. However, although similar arginine rich amphiphiles have been used to encapsulate proteins and CRISPR in RNP format [49], the capacity of RALA to deliver proteins and CRISPR machinery remains unexplored.

Here, we establish the RALA peptide as an effective novel tool for delivering CRISPR machinery into therapeutically relevant primary cells for multiple editing applications. We first characterized RALA’s capacity to complex CRISPR components in pDNA, RNP, and mRNA molecular formats into stable nanoparticles. We then used these nanoparticles to transfect primary MSCs and assess transfection efficiency, cytotoxicity, and proliferation, showing that RALA outperforms commercially available non-viral vectors while maintaining cell proliferation. Next, we effectively employed RALA to deliver pDNA and RNP to knock-in and knock-out red fluorescent protein (RFP) and luciferase (LUC) reporter genes. Finally, we used RALA-mRNA delivery to activate the expression of endogenous genes that could modulate stem cell differentiation towards different lineages. These results demonstrate that RALA enables versatile CRISPR delivery in diverse molecular formats, facilitating a wide range of gene editing approaches for effective stem cell therapies.

## 2 Materials and Methods

### 2.1 Materials

Gene synthesis and cloning were performed by Genscript, USA. pOTTC763 - pX458 with rat *ROSA26* gRNA A was a gift from Brandon Harvey (Addgene plasmid # 113161 ; http://n2t.net/addgene:113161 ; RRID:Addgene_113161) [50]. The RFP-LUC knock-in donor sequence was designed by first selecting miRFP670nano3 (RFP) and nanoluc (LUC) as small proteins with strong fluorescent and bioluminescent signals respectively [51, 52]. These sequences were separated by a *Thosea asigna* virus 2A (T2A) linker, placed under the cytomegalovirus (CMV) enhancer fused to chicken beta-actin promoter (CAG), and flanked by 1000 bp homology arms taken from the mRatBN7.2 genome on either side of the gRNA sequence. This entire sequence was constructed via de novo gene synthesis and cloned into pcDNA3-mRFP, a gift from Doug Golenbock (Addgene plasmid # 13032 ; http://n2t.net/addgene:13032 ; RRID:Addgene_13032), replacing the RFP sequence.

Plasmids for mRNA production were purchased from addgene. dSpCas9-VPR was a gift from Rasmus Bak (Addgene plasmid # 205247 ; http://n2t.net/addgene:205247 ; RRID:Addgene_205247) [53]. pGEM4Z-T7-5’UTR-EGFP-3’UTR-A64 was a gift from Christopher Grigsby & Molly Stevens (Addgene plasmid # 203348 http://n2t.net/addgene:203348 ; RRID:Addgene_203348). dCas9-VPR and eGFP plasmids were linearized using the Fast Digest Lgui and BcuI restriction enzymes respectively (Fisher scientific). Digested pDNA was gel purified using the QIAquick Gel Extraction Kit (Qiagen) and transcribed using the HiScribe® T7 Quick High Yield RNA Synthesis Kit (NEB). mRNA was purified using the Monarch® RNA Cleanup Kit (NEB). Yield and quality were assessed using a NanoDrop™ 2000 spectrophotometer (Thermo Scientific) and gel electrophoresis respectively.

Guide RNAs (gRNAs) for knocking-out the RFP-LUC reporter genes were designed by inputting the miRFP670nano3 sequence into CHOPCHOP with mRATBN7.2, CRISPR/Cas9 and knock-out specified [54]. The top scoring sequences starting with the G codon were selected for optimal compatibility with the precision gRNA synthesis kit (Thermo Fisher A29377). For gRNA targeting *TGFB3, BMP2,* or *TNMD* the transcriptional start site for each gene was identified through literature and 400 base pair (bp) sequences immediately upstream of the transcriptional start site (TSS) were downloaded from the National Center for Biotechnology Information (NCBI) genome data viewer [55, 56, 57, 58]. These sequences were then input into CHOPCHOP with mRATBN7.2, CRISPR/Cas9 and activation specified [54]. When possible, the top scoring sequences starting with a G were selected for better compatibility with the T7 promoter. Primers for each gRNA were designed (**Table 1**) and gRNAs were synthesized and purified according to the precision gRNA synthesis kit protocol.

**Table 1.**
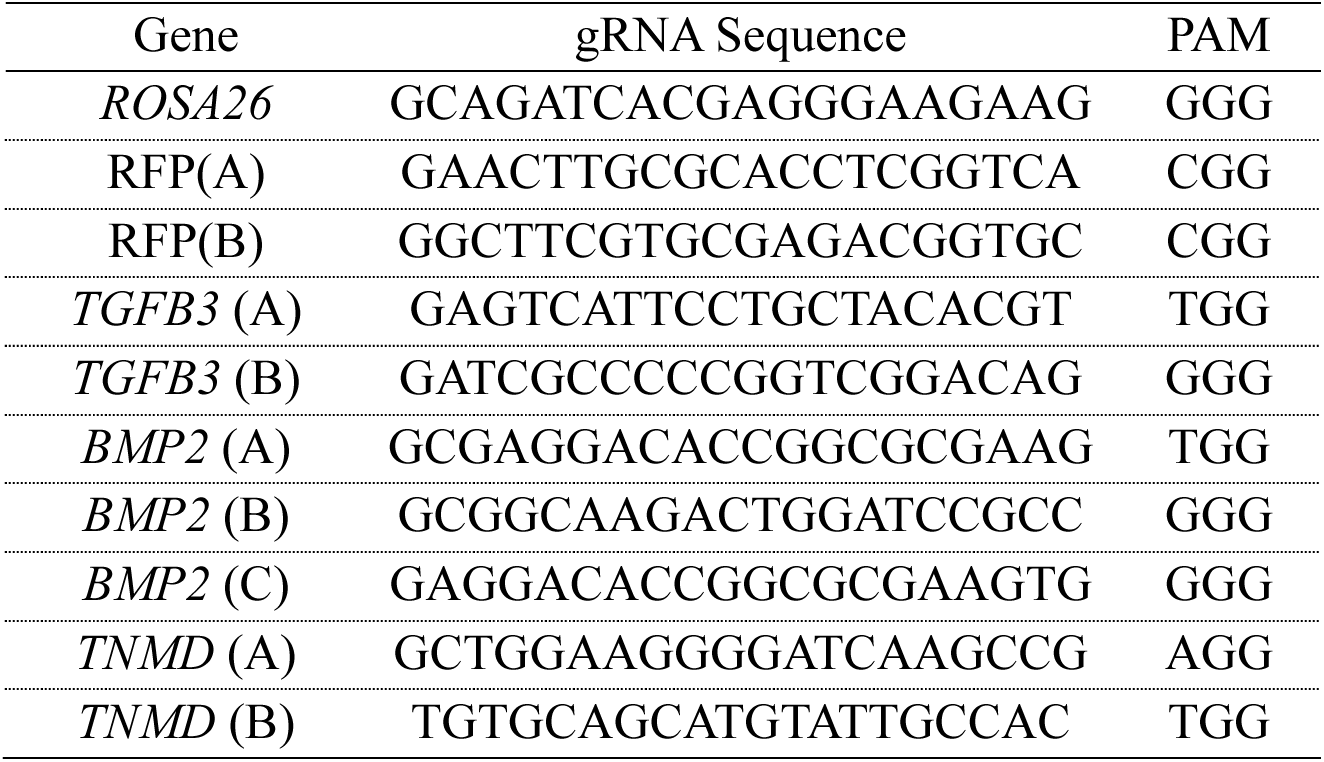
List of gRNA sequences used in this study.

TrueCut™ HiFi Cas9 Protein was purchased from Thermo Fisher. RNPs were formed by mixing Cas9 protein with gRNA at a 1:1 molar ratio for 15 minutes at room temperature. Fluorescently labeled RNPs were produced by staining gRNAs with Ulysis™ Alexa Fluor™ 594 (Invitrogen) according to the manufacturer’s protocols. Labelled gRNA was purified with P-30 gel spin columns (Biorad) which were buffer-exchanged leaving the purified gRNA in pure water.

The RALA peptide (WEARLARALARALARHLARALARALRACEA) was produced by chemical peptide synthesis at Genscript (USA) and provided as a lyophilized powder at 96.6% purity via high-performance liquid chromatography (HPLC). Lipofectamine 3000 was purchased from Thermo Fisher. Branched 25 kD polyethyleneimine (PEI) was purchased from Millipore Sigma.

### 2.2 Nanoparticle formulation and characterization

RALA nanoparticles were formed by mixing pDNA, mRNA, gRNA or RNP with the RALA peptide at the specified nitrogen:phosphate (N:P) ratios and incubating for 15 minutes. For RNP nanoparticles Cas9 and gRNA were mixed at a 1:1 molar ratio of Cas9:gRNA and incubated at room temperature for 15 minutes before adding the RALA peptide. RALA nanoparticles were characterized using dynamic light scattering (DLS) (Zetasizer nano ZS, Malvern Panalytical, UK) to measure the mean hydrodynamic diameter (size), zeta potential (charge), and polydispersity index (PDI) [59]. After complexing, 50 μL of nanoparticle mixture was diluted 1:20 vol/vol in ultrapure water and loaded into polystyrene cuvettes or folded capillary zeta cells for size and charge using automated settings.

### 2.3 MSC Isolation

Rat MSCs were harvested from male Sprague Daley rats (Charles River Laboratories) using protocols approved by the Lehigh University Institutional Animal Care and Use Committee (IACUC). Rats were euthanized and the hind leg tibia and femurs were removed and cleaned. The bone marrow from each extract was collected by flushing with warm Dulbecco’s Modified Eagle Medium (DMEM, low glucose; Gibco) via a 25-gauge needle. The resulting suspension was filtered using a 70 μm filter and seeded at 1 x 10^6^ cells/cm^2^ onto tissue culture plastic treated for cell adhesion. Cells were grown in expansion media (DMEM supplemented with 10% fetal bovine serum (FBS) (Gemini Bio) 1% penicillin/streptomycin (Thermo Fisher) for 7 days with media changes every 2-3 days. To passage cells, they were first treated with trypsin (0.25% trypsin, 0.1% disodium ethylene diaminetetraacetic acid (EDTA) in Hanks’ balanced salt solution (HBSS) (Corning) for 5 minutes. Trypsin was then inactivated with expansion media and removed by centrifugation for 5 minutes at 200x g. Cells were resuspended in expansion media and reseeded at 5.7 x 10^3^ cells/cm^2^ on tissue culture plastic treated for cell adhesion.

### 2.4 Cell culture and transfection protocol

For RALA transfections, confluent MSCs were first treated with trypsin for 5 minutes before trypsin inactivation using expansion media. Cells were pelleted by centrifugation at 200x g for 5 minutes, resuspended in phosphate buffered saline (PBS), re-pelleted, and finally resuspended in OptiMEM (Gibco). Aliquots of 1 x 10^4^ cells per cm^2^ of culture area were mixed with complexed RALA-CRISPR nanoparticles. For DNA delivery, 2 x 10^2^ ng/cm^2^ of pDNA was complexed with RALA at N:P ratios of 5, 7, or 10. For RNP delivery, 2 x 10^2^ ng/cm^2^ of Cas9 complexed to gRNA was complexed at 100x, 200x, or 500x molar ratios to RALA. Finally, for RNA delivery, 1.3 x 10^2^ ng/cm^2^ of mRNA at a 1:2 molar ratio of mRNA:gRNA was complexed with RALA at N:P ratios 10, 15, and 20. Nanoparticles were incubated with the cells for 10 minutes at 37°C before cells were moved to 6-well plates and allowed to adhere. After 4 hours, OptiMEM was aspirated, and cells were washed with PBS then cultured in 2 mL of expansion media.

Lipofectamine was used as control according to manufacturer protocols. Briefly, rat MSCs were seeded at 2.6 x 10^3^ cells/cm^2^ in 6 well plates 24 hours before transfection. For pDNA transfection, 2.08 x 10^2^ ng/cm^2^ of pDNA was mixed with the 2 μL p3000 reagent per μg DNA then combined with 3 μL/μg Lipofectamine 3000 (Invitrogen) and incubated for 15 minutes before adding over cells. For mRNA delivery, 1.3 x 10^2^ ng/cm^2^ of mRNA was directly mixed with the 3 μL/μg Lipofectamine MessengerMAX reagent (Invitrogen) and added to cells. For co-transfection with gRNA the mRNA and gRNA were mixed together before adding Lipofectamine. These nanoparticles were added directly to cells which were cultured until the necessary time points.

PEI transfections were performed as previously reported [7]. Briefly, cells were seeded at 2.6 x 10^3^ cells/cm^2^ in 6-well plates 24 hours before transfection then washed with PBS and incubated in OptiMEM 2 hours before transfection. Nanoparticles were prepared by mixing pDNA or mRNA with PEI and incubating for 30 minutes. For DNA delivery, 2.08 x 10^2^ ng/cm^2^ of pDNA was complexed with PEI at N:P of 7 [60]. For RNA delivery, 1.3 x 10^2^ ng/cm^2^ of mRNA at a 1:2 molar ratio of mRNA:gRNA was complexed with PEI at N:P 15. These particles were diluted 10:1 vol/vol in OptiMEM and 500 μL of this mixture was added to each well then diluted with 1 mL of OptiMEM after 15 minutes. Finally, OptiMEM was removed, and cells were washed with PBS and 2 mL of expansion media was added to each well 4 hours after transfection.

After transfection, all fluorescent images were taken with the BZX-800 microscope (Keyence Corporation, USA) in phase contrast mode or with the fluorescein isothiocyanate (FITC), cyanine5 (Cy5), or Texas red (TxRed) filter cubes. Samples were prepared for flow cytometry by detaching cells with 0.25% trypsin, 0.1% EDTA in HBSS for 5 minutes and resuspending in PBS. Samples were assessed using a cytoflex flow cytometer (Beckman Coulter, USA). Samples were recorded at a variable flow rate to maintain the event rate at 400 events/s. The forward scatter (FSC), side scatter (SSC), FITC, and Allophycocyanin (APC) channel gains were set to 15, 20, 1, and 25 respectively. The resulting data was gating on FSC-A and SSC-A to remove debris then gated for FSC-A and FSC-H to select the linear cluster of single cells. GFP, RFP, or Ulysis fluorophore 594 positive thresholds were defined by non-transfected rMSCs for each experiment. The transfection rate is defined as the percentage of GFP, RFP, or Ulysis Fluorophore 594 positive cells compared to the total number of single cells. The cell viability was defined as the percentage of total single cells in transfected samples versus a non-transfected control. The transfection yield is defined as the percentage of GFP, RFP, or Ulysis Fluorophore 594 positive cells compared to the total single cell count of the non-transfected control.

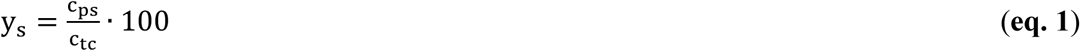

where: y_s_ = transfection yield for sample s, c_ps_ = fluorophore positive cell count, and c_tc_ = total cell count for the non-transfected control sample.

Additionally, the cell doubling time was calculated between days one and 3 using equation 2.

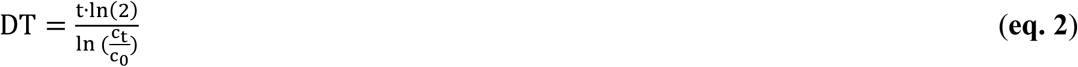

where: DT = cell doubling time, t = time in days elapsed between cell counts, c_t_ = cell count at time t, and c_0_ = cell count at initial timepoint.

### 2.5 RFP Knock-in

The knock-in of RFP was completed by co-transfecting equal amounts of the synthesized RFP-LUC donor plasmid with Addgene plasmid # 113161 containing the Cas9-T2A-GFP and *ROSA26* gRNA sequences. Cell media was treated with 250 μg/mL Geneticin (Gibco) from days 2-10 to select for cells expressing the Neomycin/Kanamycin resistance cassette within the RFP donor template. Flow cytometry was performed on days 1, 3, and 10 and fluorescent imaging was performed on days 3 and 14. On day 21 knock-in cells were purified using flow assisted cell sorting (FACS) on a SH800/MA900 cell sorter (Sony).

Genomic DNA was extracted from the RFP knock-in population and purified using the Monarch® Genomic DNA Purification Kit (New England Biotech). Whole genome sequencing (WGS) was performed by Novogene with 350 bp paired-end reads and 30x coverage (90 G of data). The *Rattus norvegicus* reference genome was downloaded from the University of California at Santa Cruz (UCSC) Genome Browser for the rnor7 [61]. This genome was then modified using reform to insert the RFP-LUC sequence between the homology arm sequences in the fourth chromosome of the genome [62]. The reformed annotation file was then used to build an index then align fastq files from WGS using the bowtie2 algorithm [63]. The read aligned binary alignment map (BAM) files will then be merged to combine all reads into one file, sorted, and indexed using SAMTOOLS [64]. Finally, CRISPResso2 is utilized to visualize the sequences and editing occurrences around the Cas9 cleavage location and confirming the knock-in sequence [65].

### 2.6 RFP knock-out

To knock-out expression of the RFP-LUC sequence, RNP was formed with Cas9 and gRNA targeting RFP. Two gRNAs were identified for RFP knock-out and are listed in **Table 1**. RNPs containing both sequences were complexed into nanoparticles with RALA at a molar ratio of 100x and transfected into purified RFP knock-in cells. As a negative control RNP targeting outside of the *ROSA26* locus (non-RFP) (gRNA sequence in **Table 1**) was transfected to ensure that loss of RFP signal was not a result of transfection. Cells were cultured for 7 days to allow for the proliferation of knock-out cells. Flow cytometry was performed on day 7 to assess the RFP signal through the APC channel. Additionally, 6.25 x 10^3^ cells/cm^2^ of each sample were passage into a 24-well plate for fluorescent imaging and quantification of luciferase expression signal. Bioluminescence was quantified by replacing media with 500 μL of 100x dilute Nano-Glo live cell substrate (Promega) in expansion media. After 30 minutes the average radiance (p/s/cm^2^/sr) of each well was quantified using a Revvity Lumina III optical *in vitro* imaging system (IVIS).

### 2.7 Gene Activation

mRNA encoding the nuclease null Cas9 fused to the tripartite transcriptional activator VPR (dCas9-VPR) was mixed with gRNA targeting either *TGFB3, BMP2, or TNMD* at a 1:2 mRNA:gRNA molar ratio and complexed with RALA at N:P ratio of 15. Total RNA was extracted 24 hours later using the Trizol reagent (Invitrogen), purified using PureLink™ RNA Mini Kit (Invitrogen), and reverse transcribed using SuperScript™ IV VILO™ Master Mix (Invitrogen). Quantitative PCR (qPCR) was performed using TaqMan™ Fast Advanced Master Mix for qPCR (Applied Biosystems) and specific gene expression assays (Applied Biosystems). TaqMan gene expression assay primers were used to monitor the expression *TGFB3, BMP2,* and *TNMD* in comparison to the *RPL13A* housekeeping control (**Table 2**). Reactions were prepared according to the manufacturers protocol in a 0.2 mL MicroAmp™ optical 96-well reaction plate with barcode (Applied Biosystems) and PCR was performed in a QuantStudio3 real time PCR system (Applied Biosystems, USA) with the ΔΔCt method. All fold changes were quantified with respect to a control transfected with no gRNA.

**Table 2.** List of TaqMan Primers used for qPCR. Gene Symbol TaqMan Assay ID.

| Gene Symbol | TaqMan Assay ID |
| --- | --- |
| *TGFB3* | Rn00565937_m1 |
| *BMP2* | Rn01449953_m1 |
| *TNMD* | Rn00574164_m1 |
| *RPL13A* | Rn00821946_g1 |

### 2.8 Statistics

Statistical analyses were performed using GraphPad Prism (version 10.2.1) software. All data was tested for normality using the Shapiro-Wilk test. For normal data with three or more groups, a one-way analysis of variance (ANOVA), or two-way ANOVA when collected on multiple days, was used with a Brown–Forsythe test and a Tukey post-test. Normal data with two groups was analyzed using a student’s t-test. Non-normal data was analyzed using the Kruskal-Wallis test. Numerical and graphical results are displayed as mean ± standard deviation, with the exception of qPCR fold-change data which is displayed as geometric mean ± geometric standard deviation. Significance was accepted at a level of p < 0.05. Sample size (n) is indicated within the corresponding figure legends.

## 3. Results

### 3.1 RALA effectively encapsulates and delivers CRISPR machinery in pDNA, mRNA, and RNP molecular cargo formats

RALA nanoparticles were formed with pDNA, mRNA, and RNP cargos encoding for CRISPR machinery at a range of relevant N:P or molar ratios. These nanoparticles were then characterized by size, charge, and transfection rate (**Fig. 1A**) [33, 39, 40]. RALA-pDNA nanoparticles at N:P ratios of 5, 7, and 10 formed stable nanoparticles with diameters consistently below 150 nm (**Fig. 1B**). Alternatively, RALA-pDNA charges positively correlated with the N:P ratio, giving zeta potentials of 16.6 ± 0.67, 21.1 ± 0.57, and 26.6 ± 3.02 mV respectively (**Fig. 1B**). This trend was also matched in PDI (Supp. Fig. 1), suggesting that higher N:P ratios of 7 and 10 form the most stable nanoparticles. Transfection of pDNA encoding for Cas9-T2A-GFP was assessed by flow cytometry and fluorescent imaging. The transfection rate (percent GFP^+^) also increased with higher N:P ratios with rates of 11.82 ± 1.53%, 18.92 ± 1.66%, and 22.52 ± 0.07% on day 1 for N:P ratios of 5, 7, and 10 respectively (**Fig. 1C-1D**, Supp. Fig. 2). However, cell viability was adversely affected at higher N:P ratios, giving N:P = 5 the highest transfection yield (percent GFP^+^ relative to non-transfected control) (**Fig. 1C**, Supp. Fig. 3).

**Figure 1.**
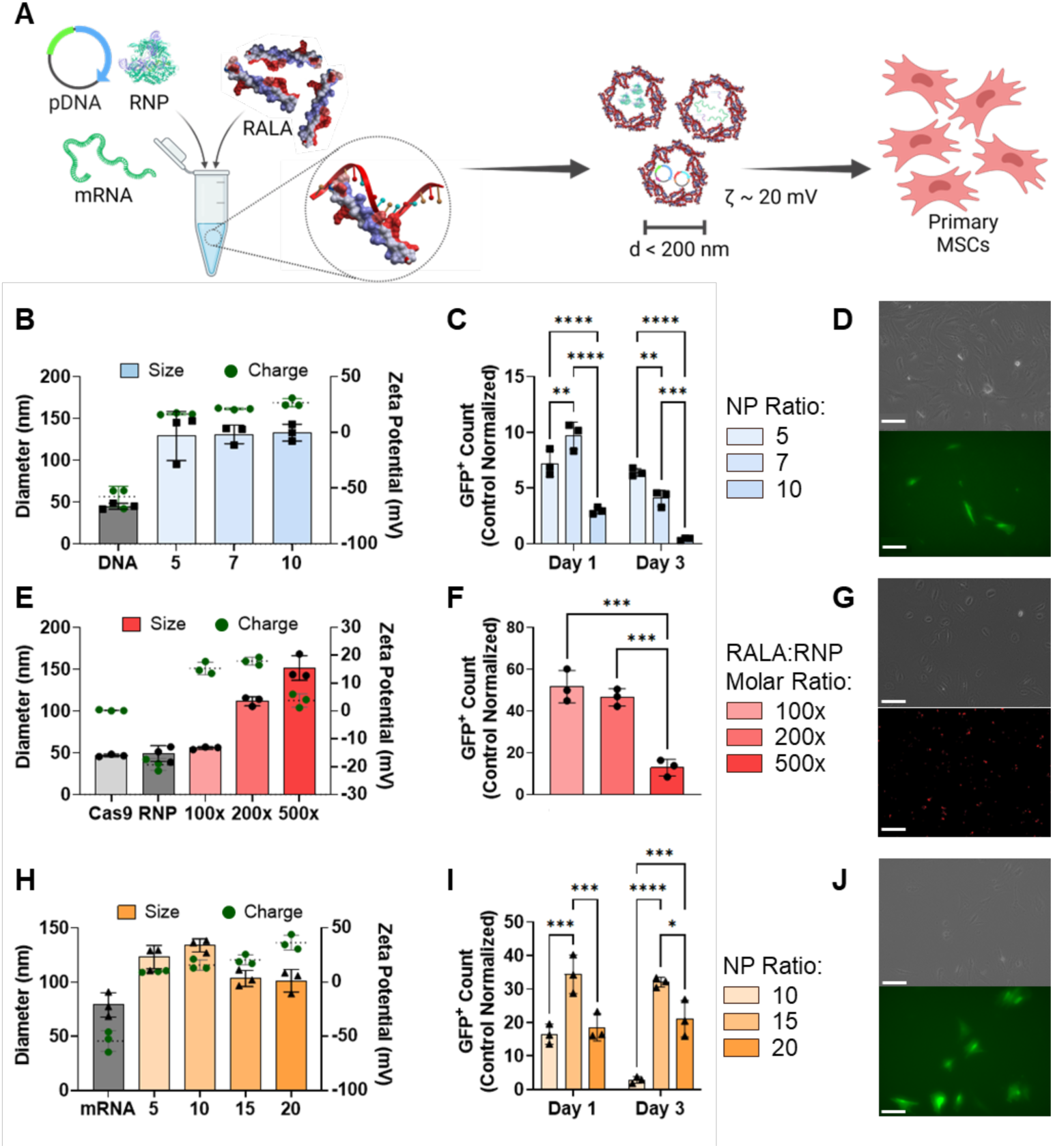
RALA encapsulates and delivers CRISPR machinery in pDNA, mRNA and RNP molecular formats into primary MSCs. **A)** Schematic of RALA-CRISPR nanoparticle formation. Individual CRISPR cargos (pDNA, mRNA, or RNP) are mixed at specific ratios with the RALA peptide resulting in positively charged nanoparticles with diameters less than 200 nm. **B)** RALA-pDNA nanoparticle size (left y axis) and zeta potential (right y axis) at different N:P and molar ratios. **C)** Transfection yield (percent transfected relative to control count) after transfection with N:P ratios of 5, 7 and 10 on days 1 and 3 after transfection, and **D)** fluorescent images of MSCs 24 hours after transfection with RALA-pDNA using N:P ratio of 5. Scale bar = 100 μm. **E)** RALA-RNP nanoparticle size (left y axis) and zeta potential (right y axis) at 100x, 200x, 500x RALA: RNP molar ratios, **F)** Transfection yield (percent transfected relative to control count) after transfection with each ratio 8 hours after transfection, and **G)** fluorescent images of 100x RALA:RNP transfection with fluorescently labelled gRNA using a Cy5 filter. Scale bar = 100 μm. **H)** RALA-mRNA nanoparticle size (left y axis) and zeta potential (right y axis) at N:P ratios of 5, 10, 15 and 20, **I)** Transfection yield (percent transfected relative to control count) after transfection with each ratio on days 1 and 3, and **J)** fluorescent images of N:P = 15 transfection one day after transfection with FITC filter. Scale bar = 100 μm. * denotes significance p<0.05 (n = 3), **p<0.01 (n = 3), ***p<0.001 (n = 3), ****p<0.0001 (n = 3).

Next, RALA-RNP nanoparticles were characterized for size, charge, and transfection capacity. Cas9 RNPs maintained a similar diameter to Cas9 alone but exhibited a negative charge demonstrating suitability for RALA complexing (**Fig. 1E**). Nanoparticle formation at RALA:RNP molar ratios of 100x and 200x resulted in diameters below 125 nm and positive charges of 15.3 ± 1.82 and 17.9 ± 1.14 mV respectively, suggesting stable nanoparticle formation. However, at the 500x molar ratio there was an increase in diameter to 151.4 ± 11.6 nm and charge reduction to 3.75 ± 1.98 mV, indicating that the nanoparticles were less stable (**Fig. 1E**). Transfection of RNPs formed with fluorescently labeled gRNA resulted in nearly 100% transfection rate regardless of molar ratio (Supp. Fig. 5). However, the 100x molar ratio group exhibited the highest cell viability giving it the highest overall transfection yield (**Fig. 1F**, Supp. Fig. 6). These results were confirmed by localized fluorescent signal of labelled gRNAs within transfected cells (**Fig. 1G**).

Finally, RALA was complexed to mRNA encoding for dCas9-VPR forming nanoparticles displaying a significant decrease (p<0.05) in particle diameter and increase in charge at N:P ratios above 15. This suggests that N:P ratios of 15 or 20 are required for forming stable complexes. (**Fig. 1H**). This trend was confirmed through transfection of eGFP mRNA, where an N:P ratio of 15 resulted in a transfection rate of 53.1 ± 1.56% compared to 23.2 ± 4.48% for N:P = 10 (Supp. Fig. 8). At an N:P ratio of 20 the transfection rate was further increased to 71.5 ± 1.42%, but a significant loss in cell viability reduced the total number of edited cells, giving N:P = 15 the highest overall transfection yield (**Fig. 1I-1J**, Supp. Fig. 8-9).

### 3.2 RALA outperforms commercial vectors for the delivery of CRISPR pDNA

We characterized RALA’s transfection efficiency of pDNA in comparison to commercial vectors Lipofectamine 3000 and polyethyleneimine (PEI). PEI was used at N:P = 7 as optimized in previous reports [7, 60], and Lipofectamine was used according to manufacturer’s protocol at a ratio of 3 μL of Lipofectamine per 1 μg of pDNA. Each of these vectors were used for Cas9-T2A-GFP pDNA transfection and flow cytometry supported by fluorescent imaging were used to assess the transfection rates and cell viability. One day post-transfection RALA achieved a transfection rate of 22.48 ± 1.41% compared to 42.26 ± 0.53% for Lipofectamine and 17.98 ± 0.61% for PEI (**Fig. 2A**). By day 3, the transfection rate decreased to 10.87 ± 1.23% for RALA whereas the Lipofectamine and PEI rates increased to 57.23 ± 2.68% and 21.21 ± 1.13% respectively (**Fig. 2A**, **Fig. 2D**). This trend was inversely correlated with cell viability in which RALA significantly outperformed both Lipofectamine and PEI by day 3 with 57.7 ± 5.2 % of cells remaining viable in comparison to 4.1 ± 0.4 % for Lipofectamine and 8.3 ± 0.4 % for PEI (**Fig. 2B**, **Fig. 2D**). This superior viability resulted in higher transfection yields of 6.3 ± 1.0 % in comparison to 2.4 ± 0.3 % and 1.8 ± 0.1 % for Lipofectamine and PEI respectively (**Fig. 2C**, **Fig. 2D**). The self-renewal capacity of transfected cells was further analyzed through quantifying cell doubling time. Cell counts from days 1 to 3 revealed that the RALA group maintained a high proliferation rate which was lost in both Lipofectamine and PEI groups (**Fig. 2E)**.

**Figure 2.**
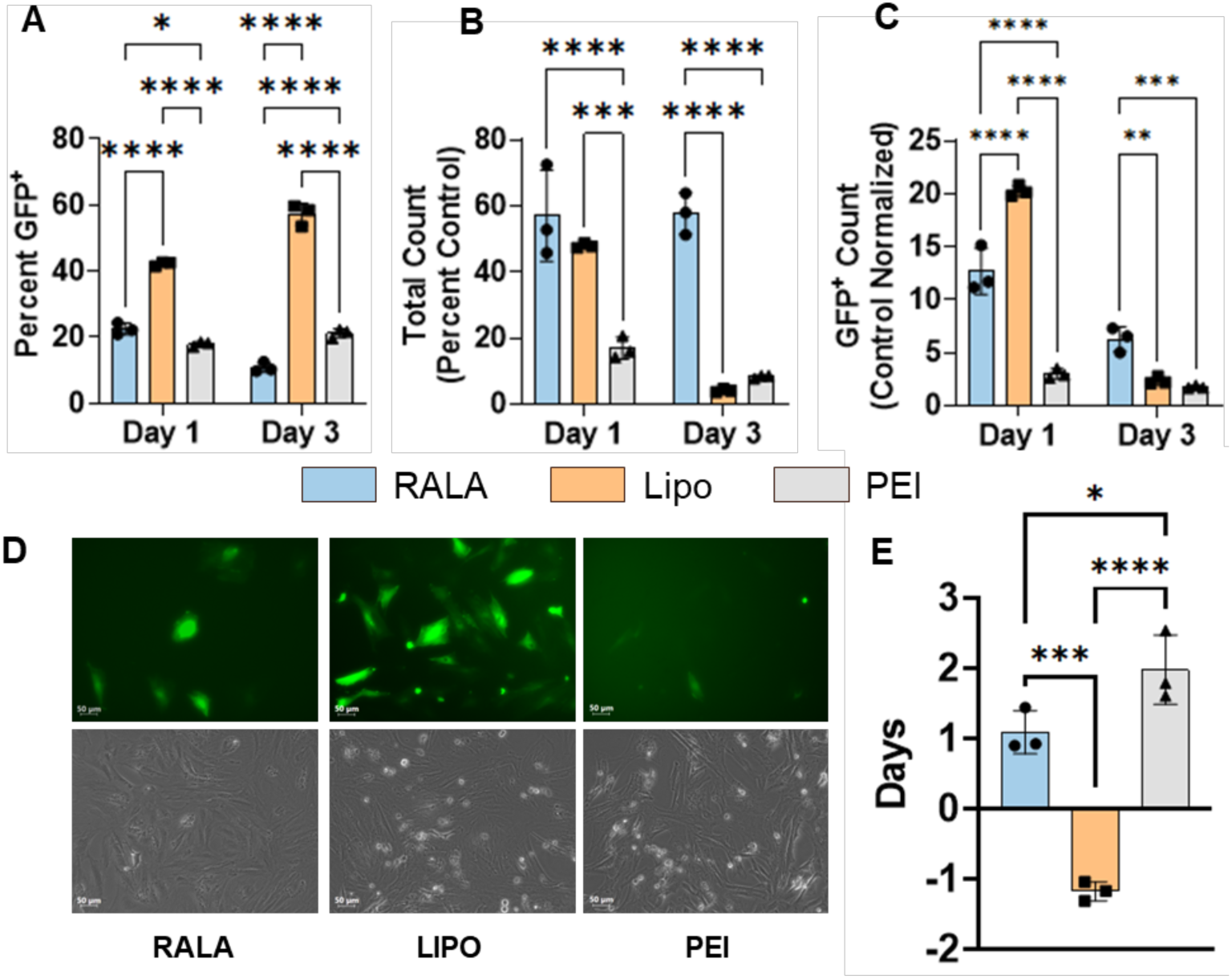
RALA outperforms commercial vectors for the delivery of pDNA encoding for CRISPR components. The transfection of pDNA encoding for Cas9-T2A-GFP using RALA, Lipofectamine 3000, or PEI was characterized via flow cytometry. **A)** Transfection rate defined as the percentage of GFP^+^ cells in the transfected population. **B)** Cell viability defined as the total cell counts as a percentage of the non-transfected control count. **C)** Transfection yield defined as the percentage of GFP^+^ cells compared to the number of non-transfected control cells. The transfection was further characterized through **D)** representative fluorescent images of each transfection and **E)** the calculation of the cell doubling time from days 1 to 3. * denotes significance p<0.05 (n = 3), **p<0.01 (n = 3), ***p<0.001 (n = 3), ****p<0.0001 (n = 3).

### 3.3 CRISPR knock-in through RALA-mediated pDNA co-delivery

After confirming RALA’s superior properties in comparison to commercial vectors, we used RALA to knock-in an RFP-LUC reporter gene into the genome of primary MSCs. We followed a strategy based on the co-delivery of two pDNAs: (1) encoding for CRISPR machinery (Cas9-T2A-GFP and gRNA) targeting the *ROSA26* locus, and (2) DNA donor sequence containing the RFP-LUC genes flanked by homology arms (**Fig. 3A**). Here, Cas9-T2A-GFP expression makes a double stranded break (DSB) within the *ROSA26* safe-harbor locus while homology arms in the donor plasmid align the RFP-LUC sequence over this break for genomic insertion (**Fig. 3A**). Flow cytometry was performed on days 1, 3, and 10 post-transfection to track the expression of each plasmid. Simultaneous expression of both plasmids was obtained in 12.15 ± 0.86% of transfected cells on day 1 (**Fig. 3B**). In the remaining cells, 9.70 ± 0.43% expressed RFP from the donor template (RFP^+^GFP^-^) compared to only 2.67 ± 0.59% expressing GFP from Cas9-T2A-GFP (RFP^-^ GFP^+^) (**Fig. 3B)**. As cells proliferated by day 3, the rate of RFP^+^GFP^+^ and RFP^+^GFP^-^ cells significantly decreased whereas a significant increase in RFP^+^GFP^-^ cells was recorded (**Fig. 3B**, Supp. Fig. 10). After 10 days of culture, GFP signal was not observed in both RFP^+^GFP^+^ and RFP^-^ GFP^+^ groups while the RFP^+^GFP^-^ percentage stayed consistent, suggesting knock-in cell proliferation (**Fig. 3B**). Fluorescent imaging at day 14 after transfection further confirmed constitutive transgene expression with the presence of RFP^+^ colonies absent of GFP expression (**Fig. 3C**). These colonies were FACS purified to a homogenous RFP-LUC population. These cells were characterized through IVIS imaging to further confirm constitutive LUC expression (Supp. Fig. 11). Finally, the genomic DNA of RFP-LUC MSCs was sequenced, confirming the presence of the RFP-LUC transgene within the genome (**Fig. 3D**).

**Figure 3.**
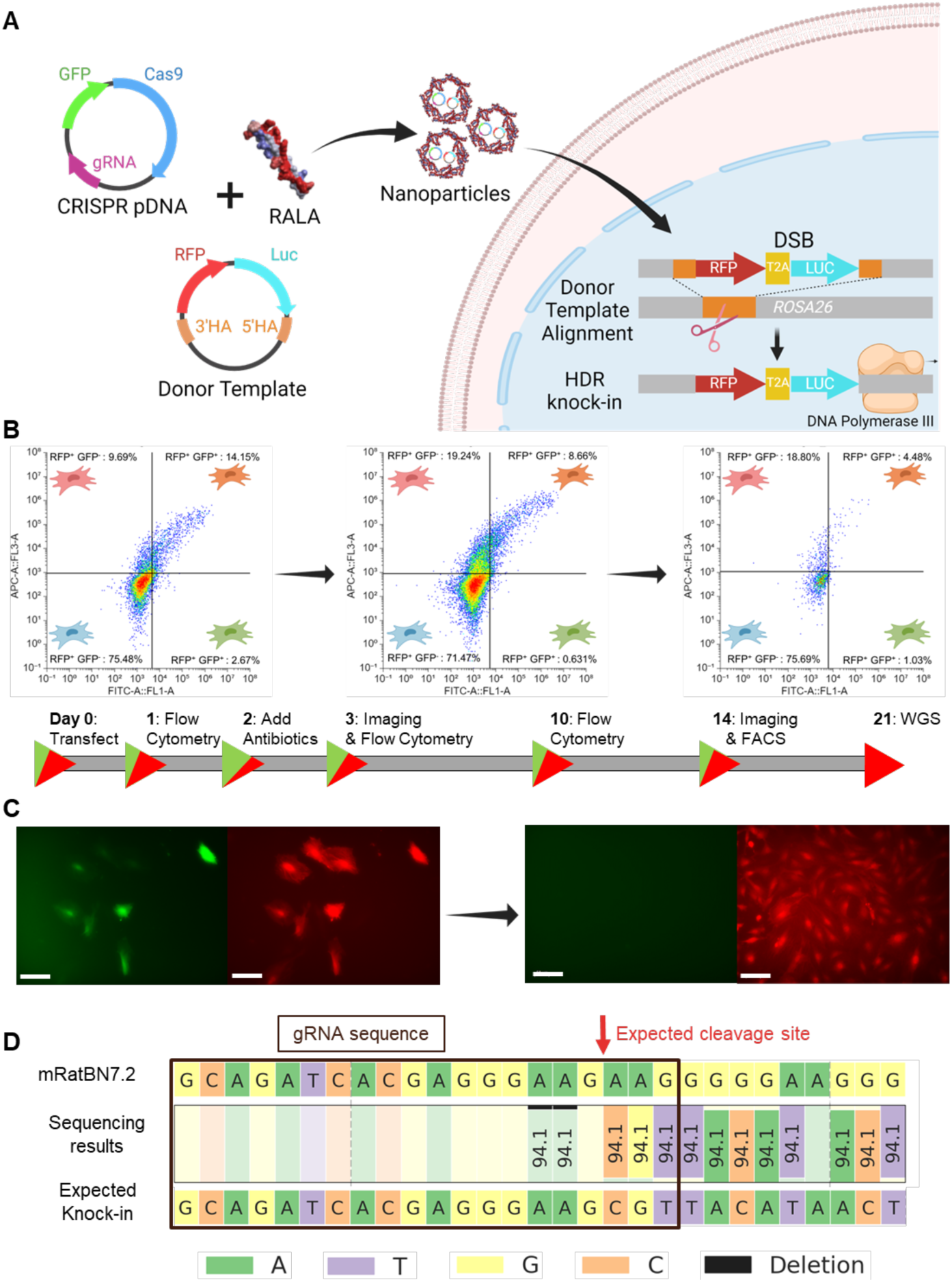
CRISPR knock-in through RALA-mediated pDNA co-delivery. **A)** Schematic of RFP-LUC knock-in strategy through pDNA co-delivery. **B)** Flow cytometry profiling of RFP knock-in cells on days 1, 3, and 10 respectively showing gating representing RFP^+^GFP^-^, RFP^+^GFP^+^, RFP^-^GFP^+^, and RFP^-^GFP^-^ cells moving clockwise from the top right. Percentages of the total population are listed. **C)** Representative fluorescent images of RFP knock-in cells taken on days 3 and 14 after transfection. Left image is FITC channel for GFP signal, right image is Cy5 channel for miRFP670nano3 imaging. Scale bar = 100 μm. **D)** WGS results at the gene knock-in location. The *Rattus norvegicus* genome assembly (top, mRatBN7.2) is compared to the expected gene knock-in sequence. The gRNA sequence used for knock-in is shown with a black box and the expected cleavage site is marked with a red arrow.

### 3.4 CRISPR knock-out through RALA-based delivery of Cas9-gRNA RNP

Next, the capacity for RALA for CRISPR knock-out (CRISPRko) through RNP delivery was assessed through the deletion of RFP expression in the previously edited RFP-LUC MSCs (**Fig. 3**). RNPs were formed with gRNA A or gRNA B targeting distinct regions within the RFP sequence. Then, the gRNAs were mixed with RALA at a 100x molar ratio as identified in section 3.1 **(Fig. 4A)**. Flow cytometry revealed a clear loss of RFP signal after one week of culture following knock-out (**Fig. 4B**). Further quantification revealed a 57.3 ± 5.5% and 14.0 ± 1.9% reduction in RFP^+^ cells after knock-out with gRNAs A and B respectively when compared to a non-RFP targeting control (**Fig. 4C**). These findings were also supported through fluorescent imaging which demonstrated a clear reduction in RFP signal following knock-out with gRNA A (**Fig. 4D**). Next, the knock-out was also characterized using an IVIS to capture the bioluminescence of each group resulting from the LUC transgene inserted downstream of RFP (**Fig. 4E**). All transfected groups experienced a significant loss in bioluminescence compared to a non-transfected control; however, no significant difference was observed between experimental and control groups (**Fig. 4F**). This result indicates that while knock-out causes a loss of RFP function, the presence of a second start codon at the LUC sequence allows continued expression.

**Figure 4.**
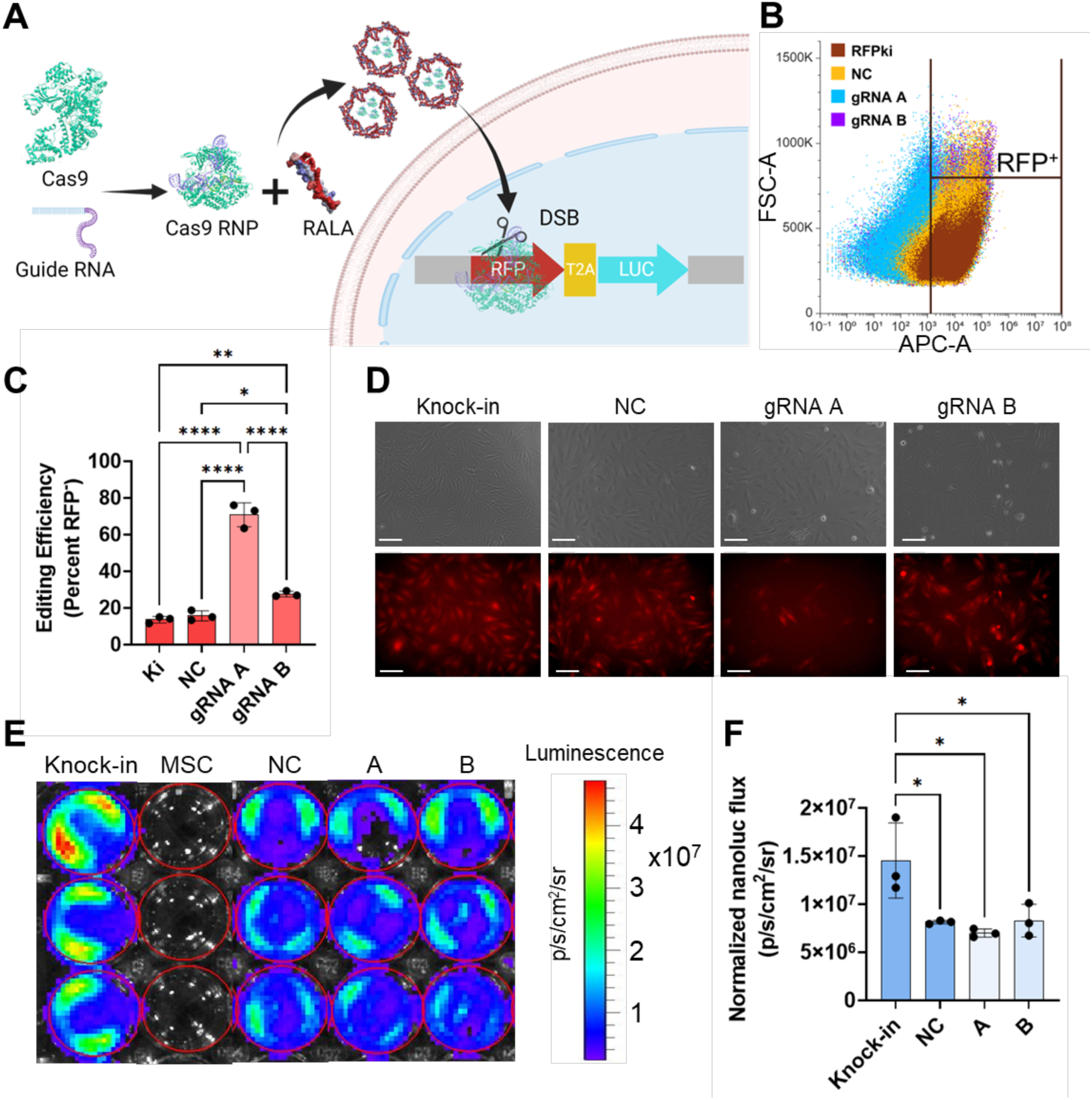
RFP knock-out using RALA-RNP nanoparticles. **A)** Schematic of gene knock-out strategy. Cas9 and gRNA targeting the RFP sequence were incubated for 15 minutes to form RNP complexes and then mixed with the RALA peptide to form nanoparticles. These nanoparticles are transfected and the RNPs are transported to the nucleus where they target the RFP gene leading to a loss of fluorescent signal. **B)** The RFP signal of each population was analyzed one week after transfection using flow cytometry to determine the rate of knock-out. **C)** The editing efficiency was defined as the percentage of RFP^-^ cells as quantified from flow cytometry results. **D)** Fluorescent images taken after passaging knock-out cells taken with Cy5 filter. Scale bar 100 μm. **E)** Average radiance (p/s/cm^2^/sr) images to visualize the bioluminescence of knock-out cells after passaging and **F)** quantification of average radiance. * Denotes significance p<0.05 (n = 3), **p<0.01 (n = 3), ***p<0.001 (n = 3), ****p<0.0001 (n = 3).

### 3.5 RALA-mRNA delivery outperforms commercial transfection reagents

In comparison to CRISPRki and CRISPRko, transcriptional regulation via CRISPR allows for precise control over gene expression without altering genomic sequences. dCas9-VPR allows activation of specific target genes depending on the chosen gRNA. However, this CRISPR tool requires efficient cellular delivery to achieve strong therapeutic effects [3]. To best accomplish this, we chose to deliver dCas9-VPR as mRNA. This allows for rapid translation in the cytoplasm in a greater number of cells in comparison to pDNA delivery [39, 42]. The capacity for RALA to effectively deliver mRNA for CRISPR transcriptional activation was determined through direct comparison to commercially available Lipofectamine MessengerMAX or PEI for the transfection of mRNA encoding for GFP. On day 1, the RALA group was outperformed by Lipofectamine in transfection rate with GFP^+^ rates of 53.13 ± 1.55% and 97.18 ± 0.22% respectively (**Fig. 5A**, **Fig. 5 D**). However, RALA achieved higher cell viability of 64.81 ± 8.50% compared to 26.0 ± 2.9 % for Lipofectamine (**Fig. 5B**, **Fig. 5D**). This elevated viability led to the RALA group significantly outperforming Lipofectamine with a transfection yield of 34.4 ± 4.7% compared to 25.8 ± 2.9% respectively (**Fig. 5C**, **Fig. 5 D**). PEI transfected cells saw a significantly lower transfection rate of 11.3 ± 1.7 % on day 1, resulting in a lower yield of 7.3 ± 1.6 % (**Fig. 5A**, **Fig. 5C**). These trends continued on day 3 when the viability of RALA-transfected cells rose to 79.73 ± 1.02% as transfected cells proliferated faster than non-transfected cells. In contrast, the viability of Lipofectamine transfected cells fell to 10.60 ± 1.14% by day 3 (**Fig. 5B**). Lipofectamine’s negative effects on cell viability were also reflected in the negative doubling time, whereas there was no difference between RALA and PEI groups (**Fig. 5E**). Overall, RALA transfections resulted in a significantly greater transfection yield of 32.0 ± 1.2 % on day 3 compared to 5.5 ± 0.7 % and 3.5 ± 0.5 % for Lipofectamine and PEI respectively.

**Figure 5.**
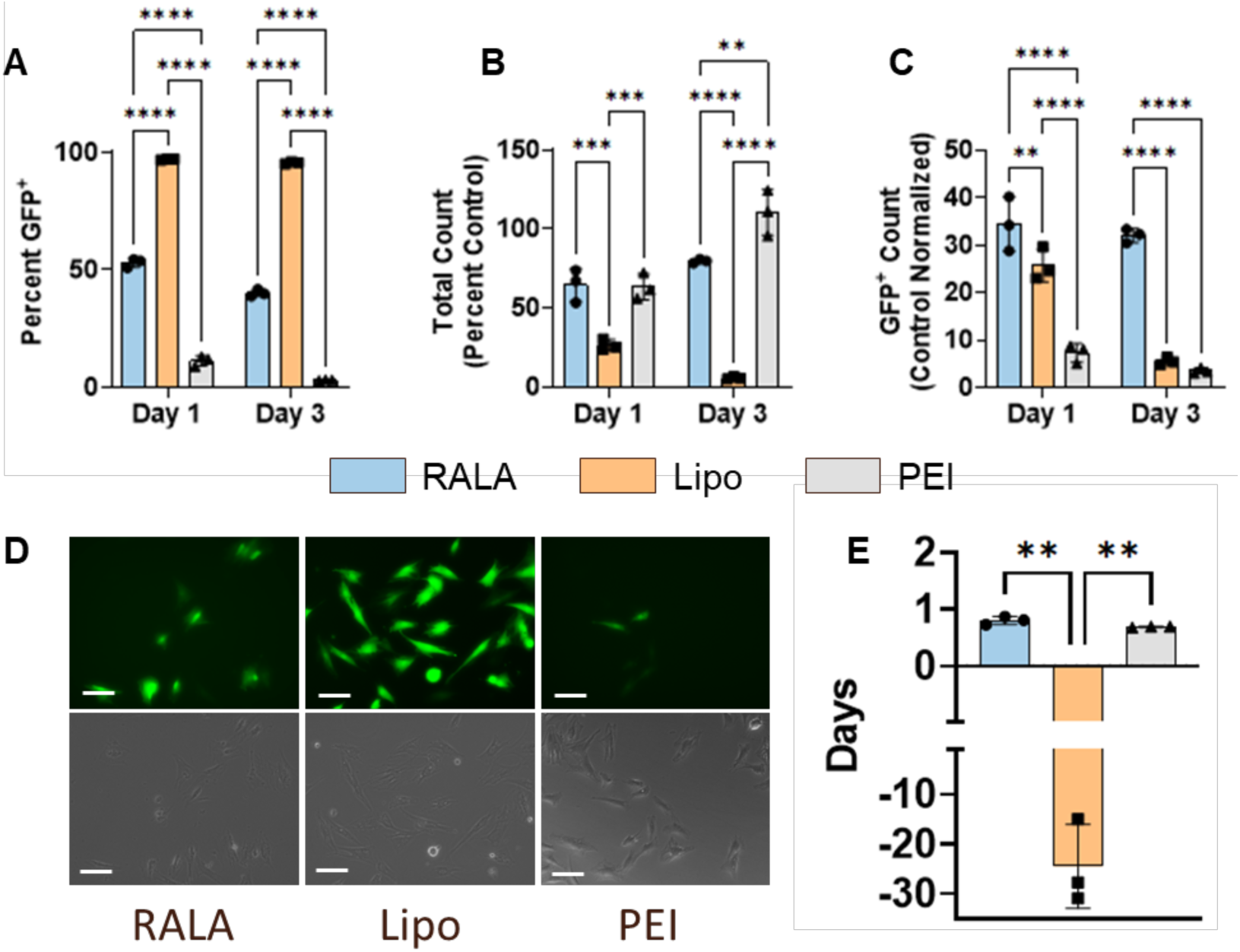
RALA mRNA delivery outperforms commercial transfection reagents Lipofectamine MessengerMAX and PEI. **A)** Transfection rate defined as the percentage of GFP^+^ cells in the transfected population at days 1 and 3 post-transfection. **B)** Cell viability defined as the total cell counts as a percentage of the non-transfected control count. **C)** Transfection yield defined as the percentage of GFP^+^ cells compared to the number of non-transfected control cells. The transfection was further characterized through **D)** representative images of each transfection and **E)** the cell doubling time from days 1 to 3. * denotes significance p<0.05 (n = 3), **p<0.01 (n = 3), ***p<0.001 (n = 3), ****p<0.0001 (n = 3).

### 3.6 RALA-mRNA delivery for CRISPR-mediated activation of gene expression

Upon confirming the suitability of RALA for mRNA delivery, mRNA encoding for dCas9-VPR was co-delivered with gRNA for CRISPR-mediated activation of gene expression (CRISPRa) (**Fig. 6A**). gRNAs were designed to target the therapeutic genes transforming growth factor β3 (*TGFB3),* bone morphogenic protein 2 *(BMP2),* and tenomodulin *(TNMD).* These are well characterized proteins which overexpression drives MSC chondrogenic, osteogenic, and tenogenic differentiation respectively (**Table 1**) [66, 67, 68]. *TGFB3* and *BMP2* were both moderately upregulated with fold changes of 1.69 ± 0.075 and 1.95 ± 0.19 respectively, while *TNMD* was highly upregulated at 25 ± 6.0-fold (**Fig. 6B-6D**). These activation fold-changes inversely correlated to the basal expression of each gene as indicated by the normalized Ct values (Supp. Fig. 12).

**Figure 6.**
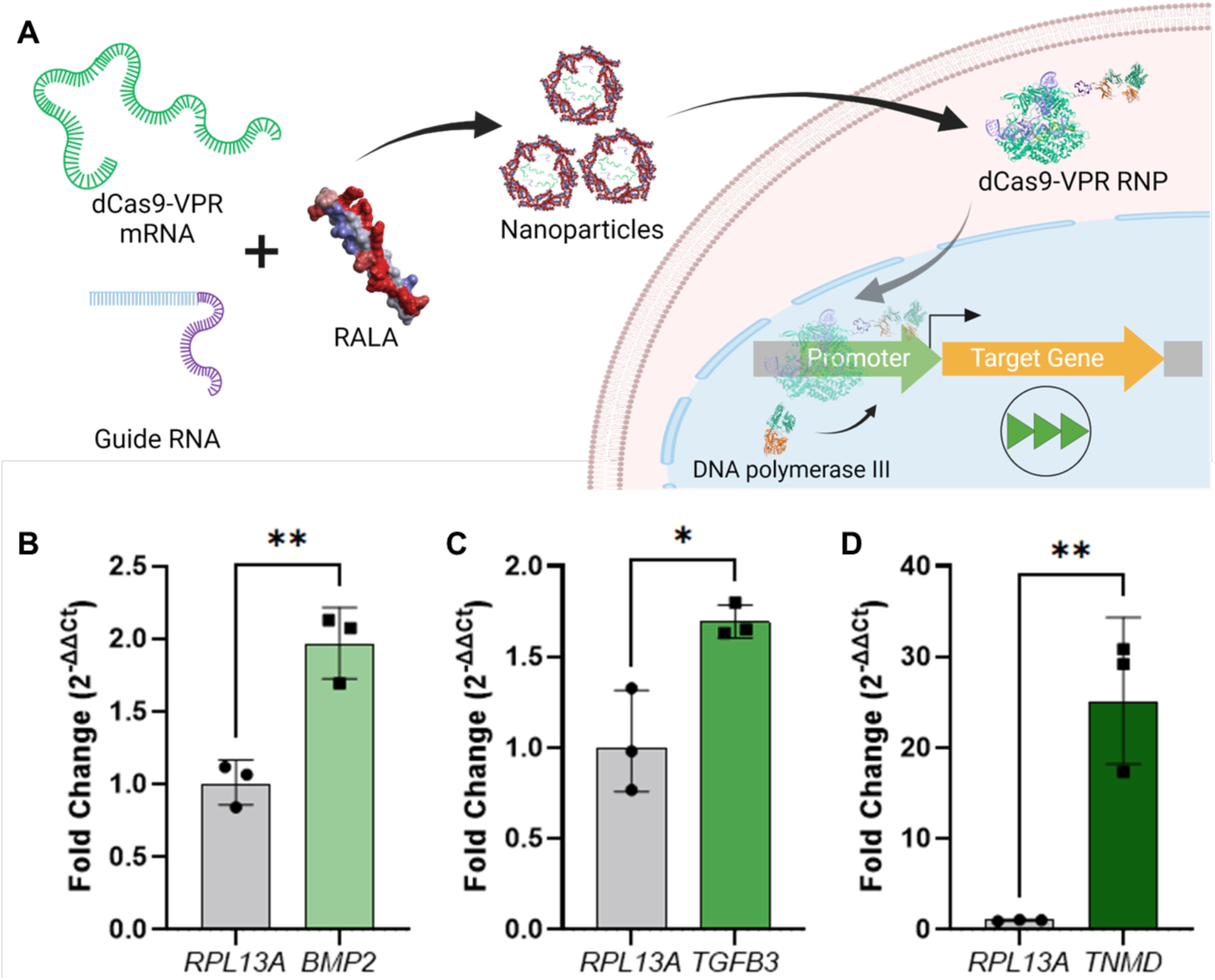
Activation of therapeutic gene expression using dCas9-VPR mRNA. **A)** Schematic of CRISPR-mediated activation of gene expression (CRISPRa) strategy followed in this study. dCas9-VPR mRNA is co-delivered with gRNA and immediately forms an RNP upon cell delivery and mRNA translation. This RNP translocates to the nucleus where it binds upstream of the target gene TSS and recruits RNA polymerase III to the promoter region to upregulate transcription. Fold changes resultant of RALA gene activation for **B)** *BMP2,* **C)** *TGFB3, and* **D)** *TNMD*. All genes are compared to a control transfected with no gRNA and the *RPL13A* housekeeping gene. * denotes significance p<0.05 (n = 3), **p<0.01 (n = 3), ***p<0.001 (n = 3), ****p<0.0001 (n = 3).

## 4. Discussion

In this study, we establish for the first time the RALA peptide as an efficient delivery vector for a wide range of CRISPR gene editing techniques in therapeutically relevant primary MSCs. Analysis of the size and charge of RALA-CRISPR nanoparticles confirmed the encapsulation of CRISPR machinery in pDNA, mRNA, and RNP formats. Flow cytometry and fluorescent imaging validated effective cellular delivery with low cytotoxicity. Each of these CRISPR formats were delivered to primary MSCs to best suit specific gene editing techniques, demonstrating the flexibility and reliability of RALA-based gene editing. The co-delivery of pDNA encoding for CRISPR machinery and the donor template induced effective knock-in and constitutive expression of an RFP-LUC transgene to generate reporter MSCs. Subsequent delivery of Cas9 RNP targeting the RFP sequence in these reporter MSCs demonstrated robust gene knock-out and loss of RFP signal. Finally, delivery of mRNA encoding for dCas9-VPR upregulated key therapeutic gene targets for MSC chondrogenesis, osteogenesis, and tenogenesis. Taken together, these results establish RALA for the efficiently delivery of CRISPR machinery in different molecular formats to perform a wide range of gene editing modalities.

The RALA cell-penetrating peptide has been used to encapsulate and deliver multiple cargos including pDNA, mRNA, micro RNAs (miRNAs), and small molecules [69, 70, 71]. However, there are no previous reports using RALA for CRISPR gene editing. In this study, RALA serves as delivery vector for diverse gene editing strategies by delivering all CRISPR molecular formats in a cell-friendly manner. RALA’s versatility is accomplished through simple electrostatic interactions between its own positively charged arginine repeats and negatively charged molecules within its cargo. However, optimizing the charge ratios between RALA and CRISPR components is essential for producing stable nanoparticles with suitable transfection capabilities. Hence, the N:P ratio between RALA and pDNA or mRNA, and the molar ratio of RALA to RNP were assessed to optimize the delivery of CRISPR machinery [33, 39, 72]. Size and charge are key indicators of nanoparticle performance, with diameters less than 200 nm and charges of 20-30 mV being optimal for cellular uptake through endocytosis [73]. We observed that all nanoparticles fit these descriptions, but the smallest diameter and highest zeta potential were achieved with N:P = 7 for pDNA, N:P = 20 for mRNA, and 100x molar ratio for RNP. To characterize the cellular delivery of these nanoparticles, pDNA and mRNA encoding for GFP or fluorescently labelled RNPs were transfected and evaluated using fluorescent imaging and flow cytometry. We analyzed the transfection rate and cell viability of each N:P ratio. Combining these two parameters we also defined a transfection yield metric consisting of the percentage of fluorescent cells compared to the number of non-transfected control cells. This metric better describes the potential of each transfection for CRISPR approaches since effective gene editing requires high numbers of edited cells without cytotoxicity [33]. We determined that lower N:P ratios of 5 and 15 for pDNA and mRNA respectively and a 100x RALA:RNP molar ratio resulted in the highest number of edited cells. This result is similar to previous reports establishing an optimal N:P ratio of 6 for pDNA delivery in MSCs [7, 39]. However, we found that a higher N:P ratio of 15 was required for mRNA compared to previous reports using an N:P ratio of 10 [42]. Additionally, we established 100x RALA:RNP as the optimal molar ratio for RNP delivery [42, 74]. These results demonstrate that RALA can effectively encapsulate and deliver all CRISPR molecular formats. This capability enables the optimization of gene editing techniques by directly comparing the editing effects of different CRISPR cargos, without using multiple delivery vectors. Additionally, due to its low cytotoxicity, RALA shows potential for complex gene editing approaches requiring the delivery of multiple components and repeated transfections [75].

CRISPRki is a powerful tool allowing the insertion of customizable transgenes via HDR for gain of function studies and the engineering of advanced cell therapies. For example, Brunger *et al.* used CRISPR to knock-in *IL1RA* and *TNFR1* genes under the cytokine-responsive *CCL2* promoter to engineer auto-regulated anti-inflammatory stem cells [76]. This work demonstrates the advantages of CRISPRki for generating synthetic gene circuits and stimuli-responsive cells. However, HDR is an inefficient process, with knock-in rates typically falling below 5% in mammalian cells [77, 78]. These rates can be improved through chemical treatments and the delivery of additional pDNAs, but these increase the risk of off-target effects, reduce cell viability, and extend *ex vivo* cell culture times [77, 78]. To increase HDR probability, CRISPR machinery can be delivered as a self-replicating pDNA, resulting in multiple gene copies over a longer time-frame [79]. Still, pDNA must be delivered to a large number of healthy and proliferative cells to allow for effective knock-in [80]. Therefore, we compared the capacity of RALA for CRISPR pDNA transfection to commercial non-viral vectors Lipofectamine 3000 and PEI. RALA outperformed PEI in transfection rate, viability, and yield, and while it had lower transfection rates than Lipofectamine, its higher cell viability resulted in better overall yields. Furthermore, cell viability was constant from days 1 to 3 after RALA transfection whereas it decreased with both Lipofectamine and PEI, demonstrating that RALA maintains the proliferative capacity of transfected cells. Enhanced cell proliferation improves HDR knock-in rates by increasing the number of cell cycles during gene editing [81]. Further, the maintenance of proliferative capacity allows for the rapid growth of edited cells which could otherwise be outcompeted by non-edited cells, limiting the overall therapeutic effect. Therefore, RALA not only delivers CRISPR machinery to a higher number of cells but also increases the likelihood that knock-in will occur during cell division.

While RALA significantly outperformed commercial vectors in this study, we observed lower cell viability post-transfection in comparison to previous reports [39, 82]. This reduced viability on day 1 is likely due to reduced cell adhesion using our suspension transfection method in comparison to previous 2D monolayer transfection [39]. Interestingly, the octa-arginine (R8) cell-penetrating peptide, containing similar arginine content to RALA, has been shown to induce strong cell-adhesion when immobilized on cell culture substrates due to charge interactions [83]. This suggests that the presence of free RALA molecules could interact with integrins in the cell membrane and decrease cell adhesion to tissue culture plastic. Despite this initial lack of cell adhesion, RALA’s capacity to transfect cells in suspension demonstrates the potential of this approach for the editing non-adherent cell types such as hematopoietic stem cells lymphocytes, and monocytes [84]. Future studies will investigate the effect of RALA on cell adhesion and the efficacy of RALA-based CRISPR delivery to non-adherent cells.

After establishing the efficacy of RALA-pDNA transfection, RALA was used for the knock-in of an RFP-LUC reporter into the *ROSA26* safe-harbor locus of primary MSCs. The knock-in and constitutive expression of these transgenes were confirmed by flow cytometry, fluorescent imaging, and WGS. The *ROSA26* locus was selected due to its constitutive expression in mammalian cells without encoding any essential genes [85]. Therefore, double stranded breaks (DSBs) can be introduced for transgene insertion without risking insertion/deletion (indel) mutations due to non-homologous end joining (NHEJ), the primary CRISPR repair pathway [86]. The continued proliferation and RFP-LUC signal from these knock-in cells demonstrates that RALA-induced gene knock-in at the *ROSA26* locus is a promising strategy for CRISPRki. Additionally, future work will target therapeutic locations to enable the quantification of gene expression [87], and the production of stimuli-responsive cells [76].

In addition to gain-of-function therapies through CRISPRki, the knock-out of specific genes also has important therapeutic implications. FDA approved CASGEVY uses CRISPRko to repress the *BCL11A* enhancer and as a result activate fetal hemoglobin for healthy red blood cell production in sickle cell anemia [88]. Such strategies benefit from the efficient delivery and transient action of CRISPR RNPs for robust editing with minimal off-target cleavage [33]. To evaluate the editing rates following RALA-RNP delivery, the reporter MSCs were treated with two RNPs targeting different locations of the RFP sequence and knock-out was assessed through flow cytometry, fluorescent imaging, and LUC quantification. A large discrepancy was observed in the efficacy of the two gRNA sequences used for this study. While gRNA A achieved a 57% reduction in RFP^+^ cells, gRNA B offered a 14% reduction. The difference in knock-out efficiency between these two gRNAs demonstrates the importance of gRNA selection and *in vitro* validation before *in vivo* studies. Yet, RALA delivery of RNPs containing gRNA A represents a reliable gene editing approach when compared to benchmark non-viral strategies. Although the previously established A5K peptide was able to achieve a higher knock-out rates of 68%, this system is designed specifically for RNP delivery and cannot deliver pDNA or other molecular formats [33]. In contrast, RALA enables a diverse range of CRISPR strategies while still accomplishing high editing rates.

Following knock-out, the bioluminescence of the edited cells was also measured. Interestingly, all transfected samples, including the negative control, had reduced bioluminescence compared to the knock-in reporter cells. These results suggest that the knock-out strategy did not enable the suppression of LUC expression. This could be due to the presence of a second start codon before LUC gene, allowing for ribosomal binding downstream of the RFP mutation for sustained LUC expression. Knock-out of both, RFP and LUC sequences could be achieved by co-delivering RNPs targeting both sequences individually. RALA has been previously established as a safe vector for repeated transfections without generating anti-RALA antibodies or over-stimulating the innate immune system [39]. Therefore, this RALA knock-in/knock-out system could serve as a powerful approach for delivering multiple transgenes into cells then selectively turning off genes once the function is no longer required.

CRISPR can also be used to transiently upregulate or inhibit target genes to induce cellular changes without permanent modification of genomic DNA. CRISPRa and CRISPR inhibition (CRISPRi) employ a Cas protein void of nuclease activity, dCas9, to deliver transcriptional machinery to specific sequences and up- or downregulate target gene expression [3]. These methods regulate the transcription of endogenous gene isoforms, thereby ensuring protein stability and reducing immunogenicity risks from foreign proteins [89]. The lack of DNA cleavage also removes indel mutation risks and improves the safety of gene editing [89]. Further, CRISPRa/i systems can be designed for multiplexed gene editing for fine control over multiple gene targets. For example, Truong *et al.* simultaneously activated *SOX9* and repressed *PPARG* to stimulate MSC chondrogenesis while preventing adipogenesis [90]. However, the viral delivery system used in this study caused inflammation, cell death, and elevated immune cell infiltration upon implantation, which limits its clinical application [90]. Conversely, RALA has been reported to have reduced immune responses *in vivo* due to transient gene delivery with reduced cytotoxicity [39, 48]. This cell-friendly delivery of CRISPRa/i components serves as a promising approach for precise transcriptomic regulation. Still, CRISPRa/i action is required in a large portion of the target cell population to achieve a significant therapeutic impact. This can be accomplished by delivering CRISPRa/i machinery as mRNA since RALA has demonstrated higher transfection rates compared to pDNA delivery [42]. Additionally, mRNA production through *in vitro* transcription is more scalable and tunable than RNP due to ease of mRNA modification, synthesis and purification [91].

The capacity of RALA to deliver mRNA encoding for eGFP was characterized in comparison to commercially available Lipofectamine MessengerMAX and PEI. RALA again outperformed PEI in all categories. In comparison to Lipofectamine, RALA resulted in a higher transfection yield due to superior cell viability. This non-cytotoxic delivery of CRISPRa machinery is essential to prevent broad transcriptomic changes that counteract or alter the effects of the desired transcriptional regulation [92]. Having established RALA for mRNA transfection, this system was then used to deliver mRNA encoding dCas9-VPR to activate therapeutically relevant genes. *TGFB3, BMP2,* and *TNMD* were selected as well-established targets for the differentiation of MSCs into chondrocytes, osteoblasts, and tenocytes respectively [66, 67, 68]. We found that the fold changes of *TGFB3* and *BMP2* were lower than *TNMD* after activation. This follows an inverse correlation to the basal expression of each gene in bone marrow MSCs [93, 94, 95]. This observation fits with previous literature suggesting that activation fold changes depend on basal gene expression [96, 97]. Konermann *et al.* demonstrated a Pearson’s correlation (r) of 0.94 (p<0.0001) between gene activation and the inverse of basal transcript levels across multiple genes [97]. Importantly, this inverse correlation suggests that activation increases the number of transcripts produced at relatively consistent rates, so the therapeutic effect is independent of the final fold change. For example, a recent study used CRISPRa to upregulate the highly expressed *VEGFA* gene in endothelial cells, only achieving ∼2-fold activation *in vitro*. Despite this low fold change, the large increase in *VEGFA* transcripts resulted in profound impacts on healing speed and quality *in vivo* as determined by wound closure, GAG deposition, and vascularization [98]. Therefore, although RALA-mediated gene activation did not achieve high fold changes for *TGFB3* and *BMP2*, it could result in greater overall therapeutic effects.

## 5. Conclusion

In conclusion, the RALA peptide is the first non-viral vector able to safely deliver all CRISPR molecular formats to primary stem cells, facilitating a wide range of CRISPR gene editing modalities. Although RALA consistently resulted in lower overall percentages of transfected cells in comparison to Lipofectamine, enhanced cell viability improved transfection yields. This higher number of healthy edited cells improves the precision and efficacy of gene editing for a more profound therapeutic impact. However, the delivery capacity of RALA could be further improved. The transient nature of RALA transfections is beneficial for the safety of gene editing as it avoids constitutive Cas9 nuclease activity but limits the duration of transcriptional regulation techniques such as CRISPRa. Future work will investigate the sustained release of RALA nanoparticles for prolonged gene activation or inhibition. Additionally, RALA lacks cell-specificity. Future work will harness the adaptability and tunability of the RALA peptide to target individual cell types for advanced gene editing approaches. Lastly, RALA transfections are performed without the presence of serum proteins or high salt content [39]. The stability and efficiency of these nanoparticles for *in vivo* conditions warrants further study. Despite these shortcomings, RALA enables the safe and effective delivery of CRISPR cargos to primary stem cells for reliable gene editing. Therefore, it is a promising delivery approach for regenerative medicine applications and the treatment of stem cell genetic disorders.

