## Supplemental data for "Versatile Cell Penetrating Peptide for Multimodal CRISPR Gene Editing in Primary Stem Cells"

Josh Graham[1], Jose Gabriel Castro[1], Lisette Werba[1], Luke Fardone[1], Kevin P. Francis[2], Anand Ramamurthi[1], Michael Layden[3], Helen O. McCarthy[4], Tomas Gonzalez-Fernandez[1*].
[1]Department of Bioengineering, Lehigh University, Bethlehem, PA, USA.
[2]Department of Orthopaedic Surgery, UCLA, Los Angeles, USA
[3]Department of Biology, Lehigh University, Bethlehem, PA, USA
[4]School of Pharmacy, Queen's University Belfast, Northern Ireland, United Kingdom

### SUPPLEMENTAL DATA

**Figures:**

Supplemental Figure 1: Polydispersity of RALA- pDNA nanoparticles ........................................1
Supplemental Figure 2. RALA -pDNA transfection efficiency at different N:P ratios....................1
Supplemental Figure 3: RALA -pDNA viability at different N:P ratios .......................................1
Supplemental Figure 4: Polydispersity of RALA- RNP nanoparticles ...........................................2
Supplemental Figure 5: RALA -RNP transfection efficiency at different N:P ratios ......................2
Supplemental Figure 6: RALA -RNP viability at different N:P ratios ..........................................2
Supplemental Figure 7: Polydispersity of RALA- mRNA nanoparticles.........................................3
Supplemental Figure 8. RALA -mRNA transfection efficiency at different N:P ratios...................3
Supplemental Figure 9: RALA -mRNA viability at different N:P ratios ......................................3
Supplemental Figure 10: Gene knock-in flow cytometry characterization................................4
Supplemental Figure 11: Gene knock-in luciferace expression characterization........................5
Supplemental Figure 12: Relative C*t* values for *BMP2, TGFB3, and TNMD* ...........................6

**Supplemental Information**

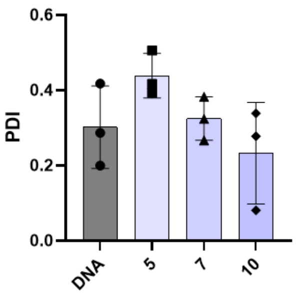


**Supplemental Figure 1.** Polydispersity index (PDI) of RALA- pDNA nanoparticles.

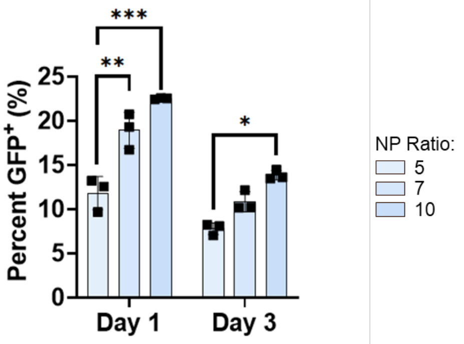


**Supplemental Figure 2.** Transfection efficiency (percent $GFP^+$) for pDNA at different N:P ratios. * denotes significance p<0.05 (n = 3), **p<0.01 (n = 3), ***p<0.001 (n = 3), ****p<0.0001 (n = 3).

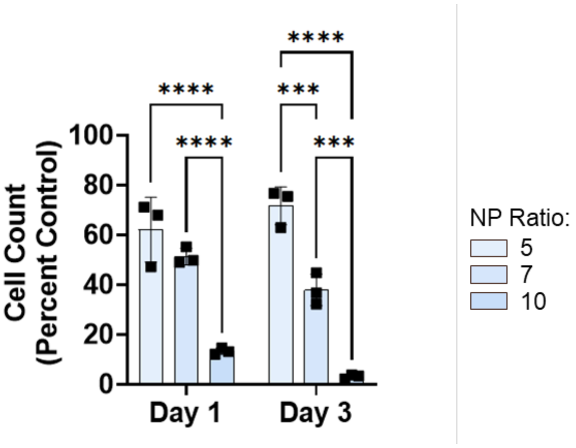


**Supplemental Figure 3.** Cell Viability (Percent cell count relative to non-transfected control) for pDNA at different N:P ratios. * denotes significance p<0.05 (n = 3), **p<0.01 (n = 3), ***p<0.001 (n = 3), ****p<0.0001 (n = 3).

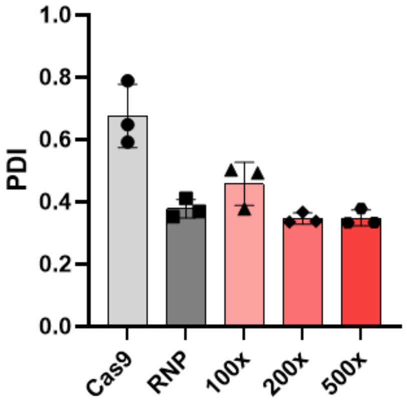


**Supplemental Figure 4.** PDI of RALA- RNP nanoparticles.

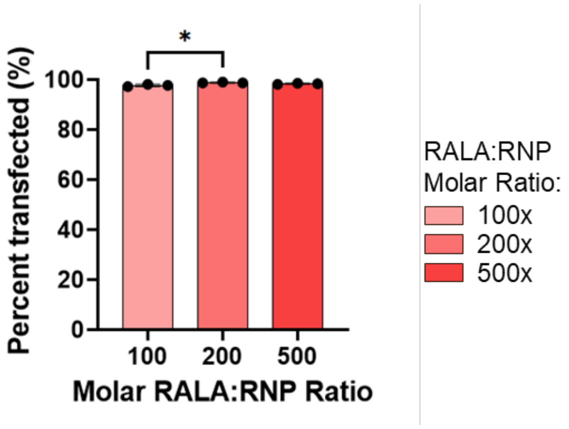


**Supplemental Figure 5**. Transfection efficiency from RNP transfections with fluorescently labeled gRNA at different molar ratios of RALA:RNP. * denotes significance p<0.05 (n = 3), **p<0.01 (n = 3), ***p<0.001 (n = 3), ****p<0.0001 (n = 3).

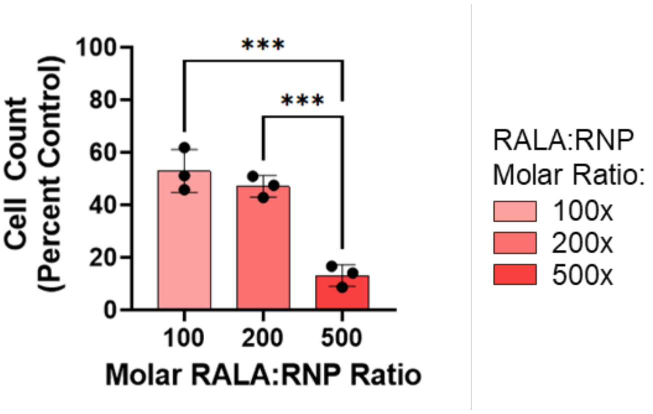


**Supplemental Figure 6**. Cell viability from RNP transfections with fluorescently labeled gRNA at different molar ratios of RALA:RNP. * denotes significance p<0.05 (n = 3), **p<0.01 (n = 3), ***p<0.001 (n = 3), ****p<0.0001 (n = 3).

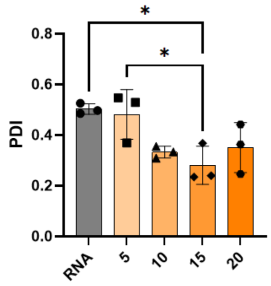


**Supplemental Figure 7.** PDI of RALA- mRNA nanoparticles. * denotes significance p<0.05 (n = 3), **p<0.01 (n = 3), ***p<0.001 (n = 3), ****p<0.0001 (n = 3).

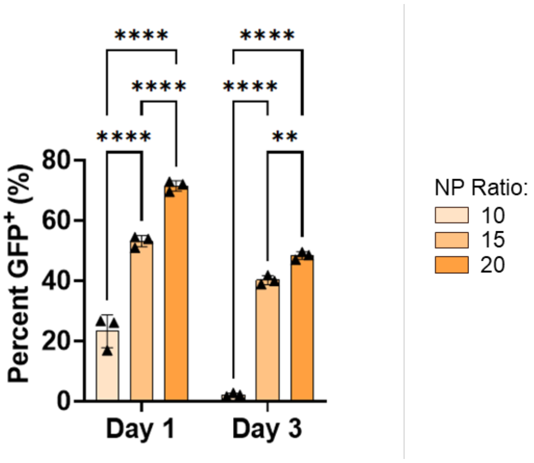


**Supplemental Figure 8.** Transfection efficiency from GFP mRNA transfections with at different N:P ratios. * denotes significance p<0.05 (n = 3), **p<0.01 (n = 3), ***p<0.001 (n = 3), ****p<0.0001 (n = 3).

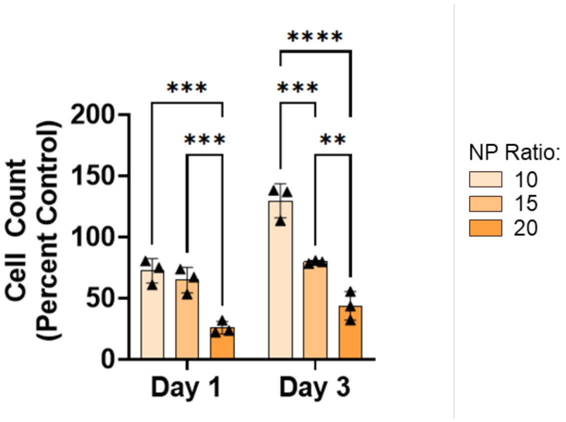


**Supplemental Figure 9.** Cell Viability from GFP mRNA transfections with at different N:P ratios. * denotes significance p<0.05 (n = 3), **p<0.01 (n = 3), ***p<0.001 (n = 3), ****p<0.0001 (n = 3).

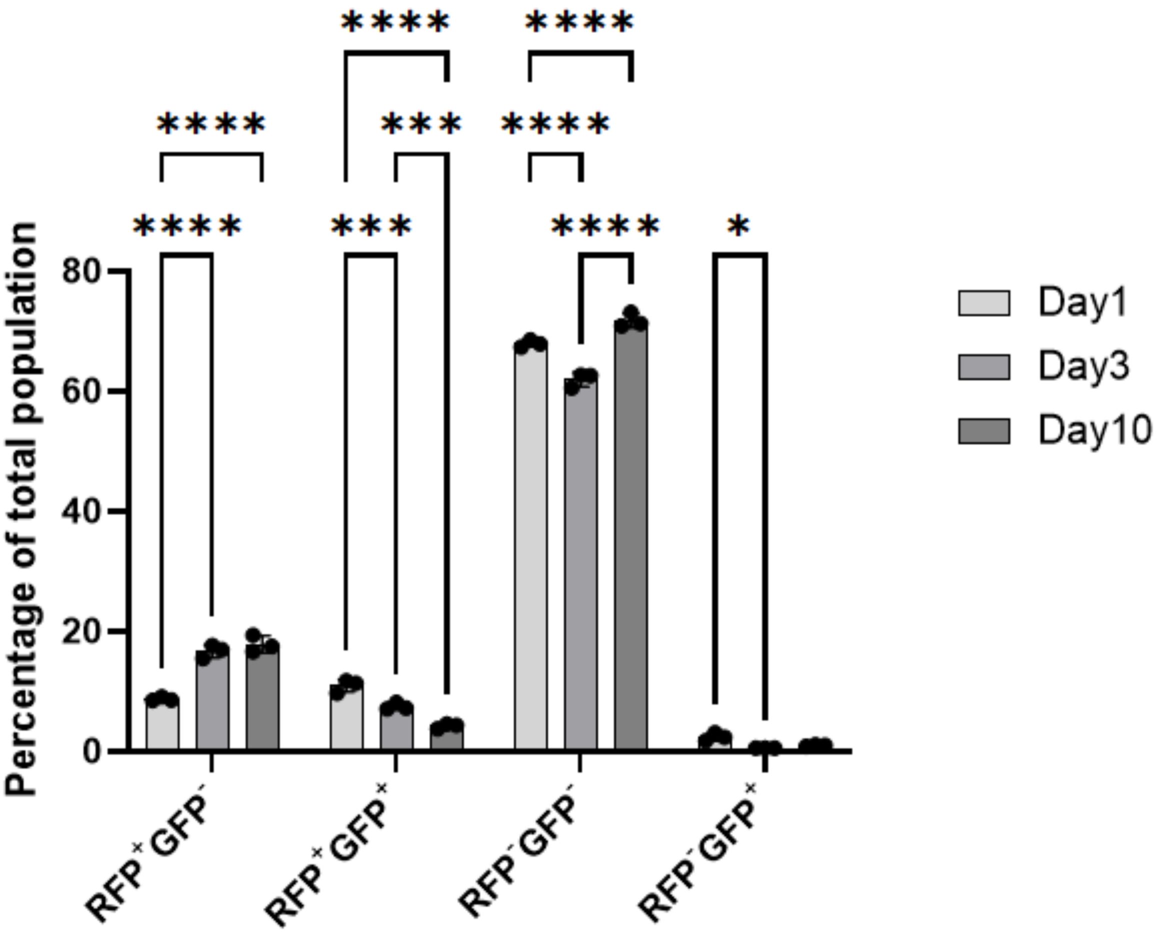


**Supplemental Figure 10.** Flow Cytometry data showing the trends in each group after co-transfection of Cas9-T2A-GFP and RFP donor sequences. Two-way anova was performed using a Tukey post-hoc test to compare each time point within each sample. * denotes significance $p<0.05$ (n = 3), **$p<0.01$ (n = 3), ***$p<0.001$ (n = 3), ****$p<0.0001$ (n = 3).

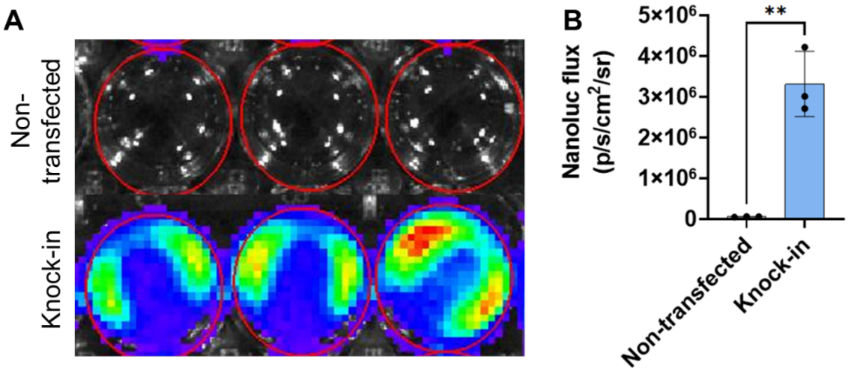


**Supplemental Figure 11**. Luciferase expression following RFP-LUC knock-in. **A)** IVIS images representing the nanoluc signal in non-transfected MSCs or RFP-LUC knock-in cells. **B)** Quantification of nanoluc flux within the region of interest (ROI) for each well further demonstrates the presence of nanoluc within knock-in cells. ** denotes significance $p<0.01$ ($n = 3$).

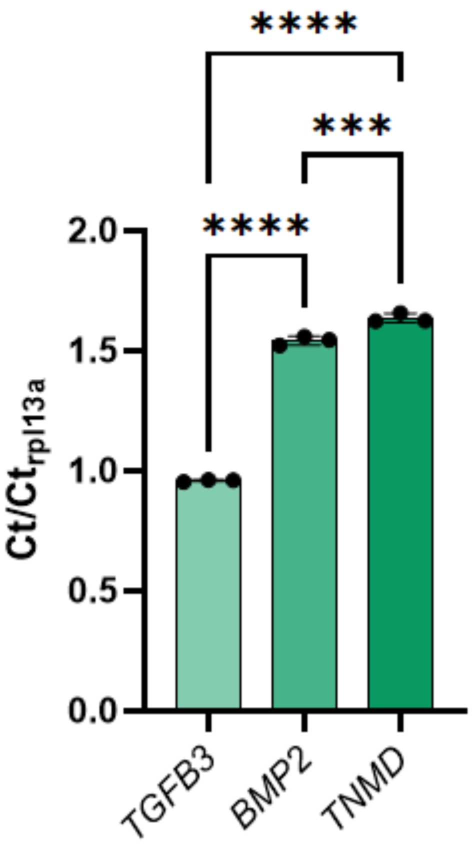


**Supplemental Figure 12.** Ct values for each gene activated using CRISPRa normalized to the Ct for the housekeeping gene (*RPL13A*). * denotes significance $p<0.05$ (n = 3), **$p<0.01$ (n = 3), ***$p<0.001$ (n = 3), ****$p<0.0001$ (n = 3).
